# Complexity of ABA signaling for stomatal development and aperture regulation

**DOI:** 10.1101/335810

**Authors:** Pirko Jalakas, Ebe Merilo, Hannes Kollist, Mikael Brosché

## Abstract

Stomata, small pores on the surfaces of leaves formed by a pair of guard cells, adapt rapidly to changes in the environment by adjusting the aperture width. As a long term response, the number of stomata is regulated during stomatal development. The hormone abscisic acid (ABA) regulates both processes. In ABA mediated guard cell signaling the protein kinase OPEN STOMATA1 (OST1) has a central role, as stomatal closure in the *ost1* mutant is impaired in response to ABA and to different environmental stimuli. We aimed to dissect the contribution of different ABA-related regulatory mechanisms in determining stomatal conductance, a combination of stomatal density and aperture width, and crossed the *ost1* mutant with mutants that either decreased (*aba3*) or increased (*cyp707a1/a3*) the concentration of ABA in plants. The double mutant *ost1 aba3* had higher stomatal conductance than either parent due to a combination of increased stomatal aperture width and higher stomatal density. In the triple mutant *ost1 cyp707a1/a3* stomatal conductance was significantly lower compared to *ost1-3* due to lower stomatal density. Further characterization of the single, double and triple mutants showed that responses to treatments that lead to stomatal closure were impaired in *ost1* as well as *ost1 aba3* and *ost1 cyp707a1/a3* mutants, supporting a critical role for OST1 in stomatal aperture regulation. Based on our results, we suggest that there are two signaling pathways to regulate water flux from leaves i.e. stomatal conductance: an ABA-dependent pathway that determines stomatal density independent of OST1; and an OST1-dependent pathway that regulates rapid changes in stomatal aperture.

## Introduction

Stomata, formed by a pair of guard cells, are small pores responsible for gas exchange in leaves. They allow CO_2_ uptake for photosynthesis, with the accompanying loss of water. In addition, air pollutants and some pathogens enter the plant through stomata. Hence, accurate adjustment of the stomatal pore is required for the plant to successfully thrive in a changing environment. Not only the width of the stomatal aperture, but also the number of stomata is regulated by environmental signals and influences plant gas exchange (Hetherington and Woodward, 2003). Several studies have shown that doubling of ambient CO_2_ concentration leads to a reduction in stomatal density in different plant species and *Arabidopsis* accessions (Woodward and Kelly, 1995; Woodward et al., 2002). In contrast, higher light intensity significantly increases stomatal density (Casson et al., 2009). The main difference between the adjustment of the stomatal aperture versus the number of stomata is the scale of time, stomatal aperture can change in minutes, whereas changes in stomatal density are fixed during leaf development.

The plant hormone abscisic acid (ABA) plays a central role in the regulation of guard cell function (Kim et al., 2010; Kollist et al., 2014). ABA-induced stomatal closure is initiated by binding of the hormone to PYR/RCAR receptors that leads to the inactivation of type 2C protein phosphatases (PP2Cs), which in turn releases SNF-related protein kinases (SnRK2s) such as OPEN STOMATA1 (OST1) to activate guard cell ion channels including SLOW ANION CHANNEL1 (SLAC1). This leads to the efflux of anions, followed by potassium and water efflux and stomatal closure (Kim et al., 2010; Kollist et al., 2014).

As ABA is the central regulatory molecule of stomatal function, fine-tuning of ABA levels and signaling is of utmost importance during acclimation to abiotic stress, e.g. drought. Guard cell ABA levels are regulated by de novo biosynthesis, catabolism, and transport from other plant tissues (Nambara and Marion-Poll, 2005; Merilo et al., 2015; Merilo et al., 2018). OST1 appears to have a critical role in ABA signaling. Stomatal closure induced by ABA or environmental factors is strongly impaired in *ost1* mutants (Mustilli et al., 2002; Merilo et al., 2013). In addition to stomatal regulation, ABA affects stomatal development, which is also controlled by environmental factors such as light and the level of CO_2_ (Casson and Hetherington, 2010; Chater et al., 2015). ABA-deficient mutants have increased stomatal densities compared to wild-type (Tanaka et al., 2013; Chater et al., 2015), whereas the ABA over-accumulating *cyp707a1/a3* double mutant had significantly lower stomatal density than wild-type (Tanaka et al., 2013), supporting the role of ABA in stomatal development. ABA3 encodes a molybdenum cofactor sulfurase required by an abscisic aldehyde oxidase to catalyze the conversion of abscisic aldehyde to ABA; its expression level increases in response to drought and ABA treatment (Xiong et al., 2001). In non-stressed conditions, the concentration of leaf ABA is approximately 45% of wild-type ABA in *aba3-1* (Merilo et al., 2018). The predominant ABA catabolic pathway, ABA 8’-hydroxylation, is mediated by four members of the CYP707A gene family and their transcription levels increase in response to salt and drought stress as well as ABA (Saito et al., 2004). CYP707A1 and CYP707A3 are important for post-germination growth, since seedling growth by exogenous ABA was inhibited more effectively in *cyp707a1* and *cyp707a3* mutants and was more pronounced in the double mutant that also contained higher concentration of ABA compared to the single mutants (Okamoto et al., 2006). Both *cyp707a1* and *cyp707a3* loss-of-function mutants showed reduced stomatal conductance, which was more pronounced in *cyp707a3* (Merilo et al., 2013).

Here we report that while OST1 is required in rapid stomatal responses to several environmental conditions: reduced air humidity, darkness, elevated CO_2_ concentration and exogenous ABA, there are alternative ABA signaling pathways in guard cells that contribute to stomatal development. These two pathways, OST1-dependent signaling that regulates stomatal aperture width and OST1–independent signaling that regulates stomatal density, coordinate the overall water flux through stomata.

## Materials and Methods

### Plant material, growth and gas-exchange measurements

Col-0, *aba3-1* and *ost1-3* (*srk2e*, SALK_008068) were from the European Arabidopsis Stock Centre (www.arabidopsis.info). The *cyp707a1 cyp707a3* double mutant was a gift from Eiji Nambara (Okamoto et al., 2006). Double mutants and other crosses were made through standard techniques and genotyped with PCR-based markers.

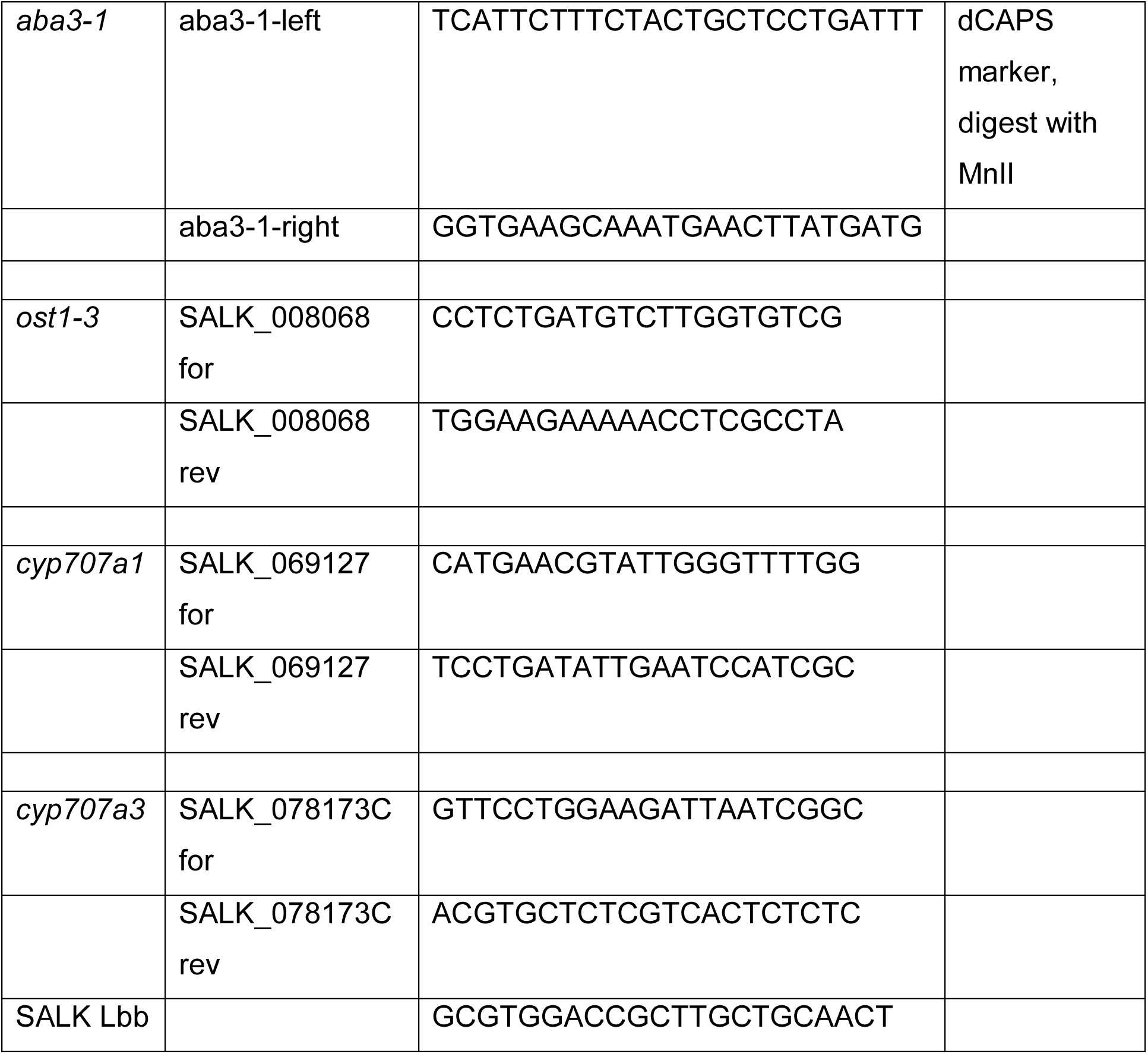

Plants for gas-exchange measurements were sown into 2:1 (v:v) peat:vermiculite mixture and grown through a hole in a glass plate covering the pot as described in Kollist et al. (2007). Plants were grown in growth chambers (AR-66LX, Percival Scientific, IA, USA and Snijders Scientific, Drogenbos, Belgia) with 12 h photoperiod, 23/18 °C day/night temperature, 150 μmol m^-2^ s^-1^ light and 70% relative humidity. Plants were 24-30 days old during gas-exchange experiments.

Stomatal conductance of intact plants was measured using a rapid-response gas-exchange measurement device similar to the one described by Kollist et al. (2007) consisting of eight thermostated flow-through whole-rosette cuvettes. Plants were inserted into the measurement chambers and after stomatal conductance had stabilized, the following stimuli were applied: reduction in air humidity (decrease from 65-75% to 30-40%), darkness (decrease from 150 μmol m^-2^ s^-1^ to 0 light), CO_2_ (increase from 400 ppm to 800 ppm) and spraying plants with 0 µM, 5 µM or 50 µM ABA solution with 0.012% Silwet L-77 (Duchefa) and 0.05% ethanol. ABA-induced stomatal closure experiments were carried out as described previously (Merilo et al., 2018). Initial changes in stomatal conductance were calculated as gs18-gs0, where gs0 is the pretreatment stomatal conductance and gs18 is the value of stomatal conductance 18 min after factor application; 16min in case of ABA spraying.

### Measurement of stomatal aperture and density

Epidermal peels were stripped from four-week-old plants grown in growth chambers as described above and incubated in resting buffer (containing 10 mM MES-KOH pH6.2) for 2.5 hours. Images of stomata were taken with a Zeiss Axio Examiner.D1 microscope. Images were taken of 15 stomata per leaf and averaged to characterize the stomatal aperture of each plant. Six plants per genotype were analyzed. Stomatal aperture width was measured using the image processing software ImageJ 1.51k (National Institutes of Health, USA).

For stomatal density (SD) measurements, leaves of 5 weeks-old plants grown as described above, one leaf per plant, were excised and the abaxial side was covered with dental resin (Xantropen VL Plus, Heraeus Kulzer, Germany). The hardened resin impressions were covered with transparent nail varnish. The dried nail varnish imprints were attached to a microscope glass slide with a transparent tape and images were taken with a Zeiss SteREO Discovery.V20 stereomicroscope. 24 plants per genotype were analyzed. SD was determined from an image with an area of ∼0.12 mm^2^, taken from the middle of the leaf, close to the middle vein and calculated as: SD = number of stomata/area of the image

### Statistical analysis

One-way ANOVA was used to compare the effect of genotype on the values of stomatal conductance, aperture, density and initial change in stomatal conductance. Comparisons between individual means were done with Tukey or Tukey unequal N HSD *post hoc* tests as indicated in figure legends. Stomatal conductance values before and after application of ABA were compared by repeated measures ANOVA with Tukey *post hoc* test. All effects were considered significant at p<0.05. Statistical analyses were performed with Statistica, version 7.1 (StatSoft Inc., Tulsa, OK, USA).

### Accession Numbers

ABA3 - AT1G16540; OST1 - AT4G33950; CYP707A1 - AT4G19230; CYP707A3 - AT5G45340.

## Results

We crossed *ost1-3* into an ABA biosynthesis mutant (*aba3-1*) and to *cyp707a1 cyp707a3* (here abbreviated as *cyp707a1/a3*) that lacks two proteins involved in ABA catabolism. By doing so, we generated plants where strong ABA-insensitivity caused by impaired OST1 was combined with defective ABA biosynthesis or breakdown (Fig. 1). Steady-state stomatal conductance and rapid stomatal responses to various closure-inducing stimuli were measured in intact plants with a custom-made gas-exchange device as described before (Kollist et al., 2007). Our results showed that the *aba3-1* mutant displayed higher stomatal conductance, whereas *cyp707a1/a3* had reduced stomatal conductance compared to Col-0 wild-type (Fig. 2A), as can be expected on the basis of the ABA concentrations in these plants (Okamoto et al., 2006; Merilo et al., 2018). The double mutant *ost1 aba3* had higher stomatal conductance than either parent (Fig. 2A) and the triple mutant *ost1 cyp707a1/a3* displayed lower stomatal conductance than the single *ost1-3* (Fig. 2A).

**Figure 1.**
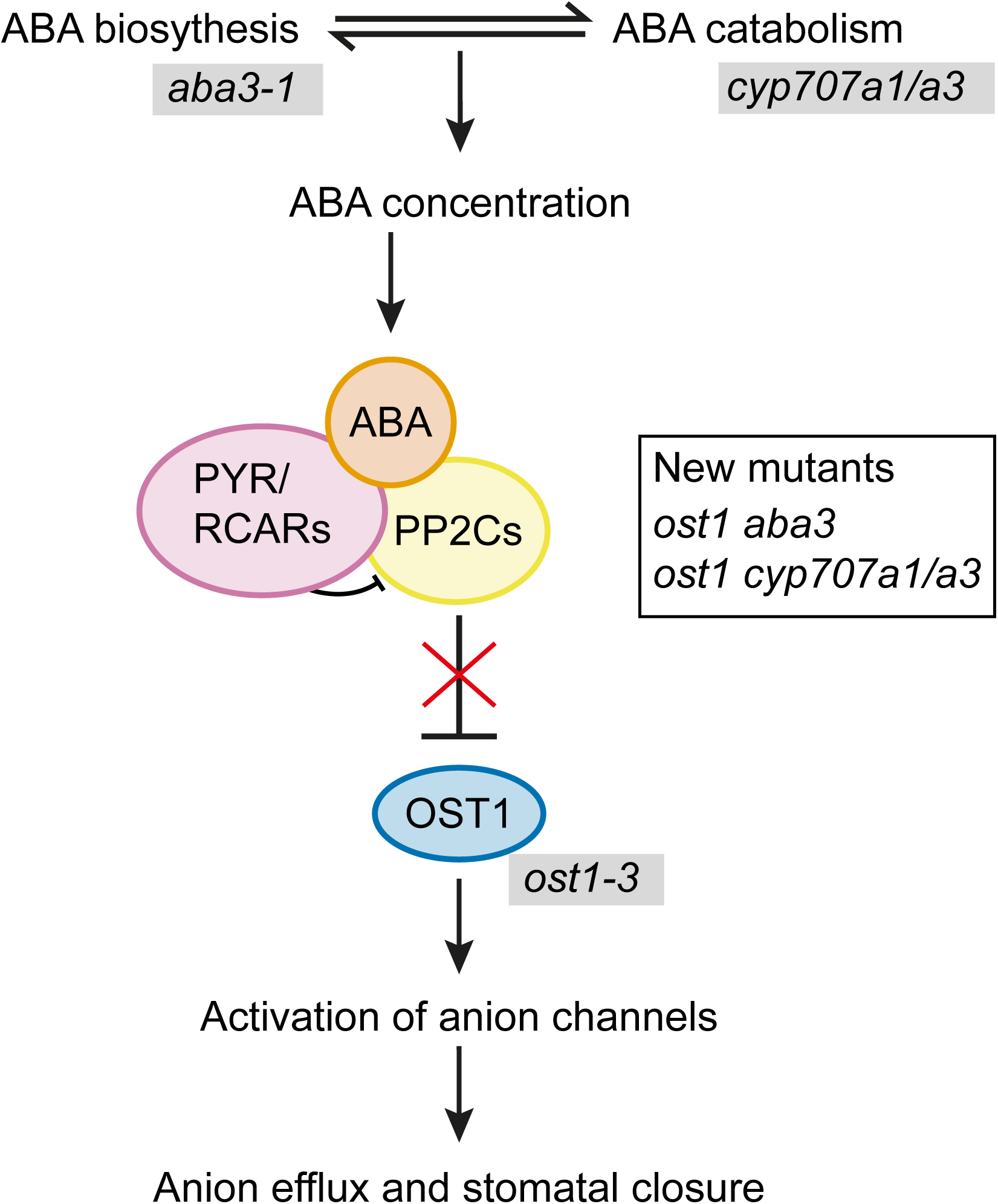
Schematic overview of ABA concentration determined by ABA biosynthesis and catabolism, followed by the core components in ABA signalling leading to stomatal closure. Mutants used in this study are indicated in grey background. New double and triple mutants generated for this study are *ost1 aba3* and *ost1 cyp707a1/a3.*

**Figure 2.**
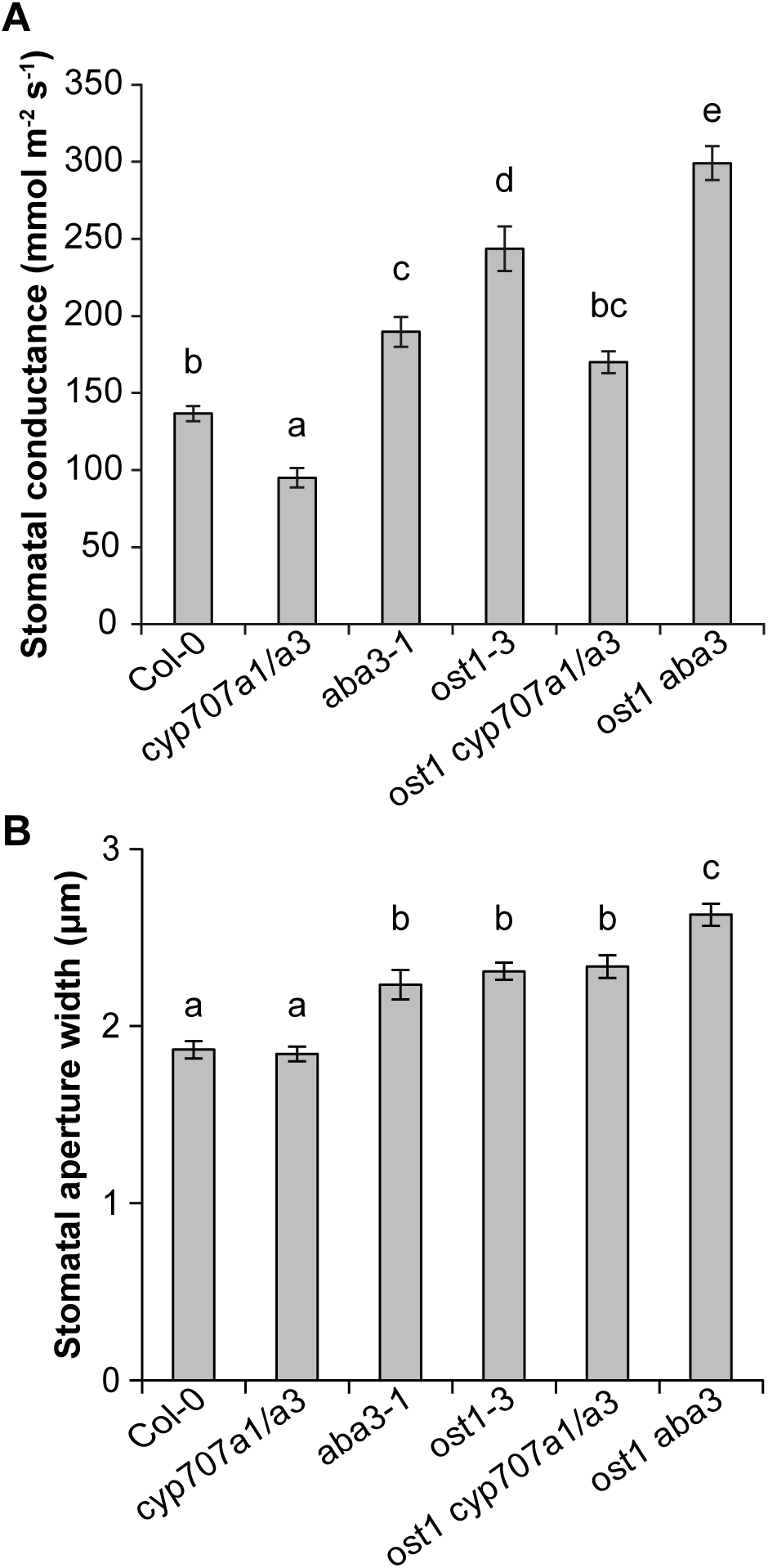
(A) Whole-plant steady-state stomatal conductance (gs) of three- to four-week-old plants. The ABA biosynthesis mutant *aba3-1* and catabolism double mutant *cyp707a1/a3* were crossed to *ost1-3* to genetically reduce or increase the ABA concentration in *ost1-3* background. Letters denote statistically significant differences between lines (ANOVA with Tukey unequal N HSD *post hoc* test, p < 0.05; n=8-13). (B) Stomatal aperture measured on epidermal peels of four-week-old plants. Letters denote statistically significant differences between lines (ANOVA with Tukey *post hoc* test, p < 0.05; n=6).

As altered stomatal conductance can result from a change in stomatal aperture width or in stomatal density, we next determined the stomatal apertures of the mutants. There were no differences in aperture widths between *cyp707a1/a3* and wild-type (Fig. 2B). Compared to wild-type, stomata of *aba3-1* and *ost1-3* single mutants had significantly wider apertures (Fig. 2B). Aperture of *ost1 cyp707a1/a3* was similar to *ost1-3*, whereas *ost1 aba3* had significantly wider aperture compared to the single mutants (Fig. 2B). These results suggest that ABA-deficiency leads to wider stomatal apertures, whereas over-accumulation of ABA seems to have no effect on aperture width. However, *cyp707a1/a3* (compared to Col-0) and *ost1 cyp707a1/a3* (compared to *ost1*) showed differences in stomatal conductance but not in aperture widths, indicating that some other trait besides aperture is involved in determining stomatal conductance.

In order to test whether the differences in stomatal conductance in the studied mutants were associated with altered stomatal density, we measured stomatal density using leaf impressions. Consistent with already published results (Tanaka et al., 2013; Chater et al., 2015), *aba3-1* had higher and *cyp707a1/a3* lower stomatal density compared to wild-type in our experiment (Figs. 3A-B). The stomatal density of *ost1-3* was similar to wild-type, but through genetically altering the ABA concentration in the *ost1-3* mutant, we could affect the stomatal development. Compared to the single *ost1-3* mutant, *ost1 aba3* and *ost1 cyp707a1/a3* had significantly higher or lower stomatal density, respectively (Figs. 3A-B). Taken together, stomatal conductance, aperture and density results show that *ost1-3* has higher stomatal conductance due to more open stomata (Figs. 2-3; Mustilli et al., 2002). However, ABA concentration is a crucial signal for stomatal development, which was apparently regulated by an OST1-independent mechanism.

**Figure 3.**
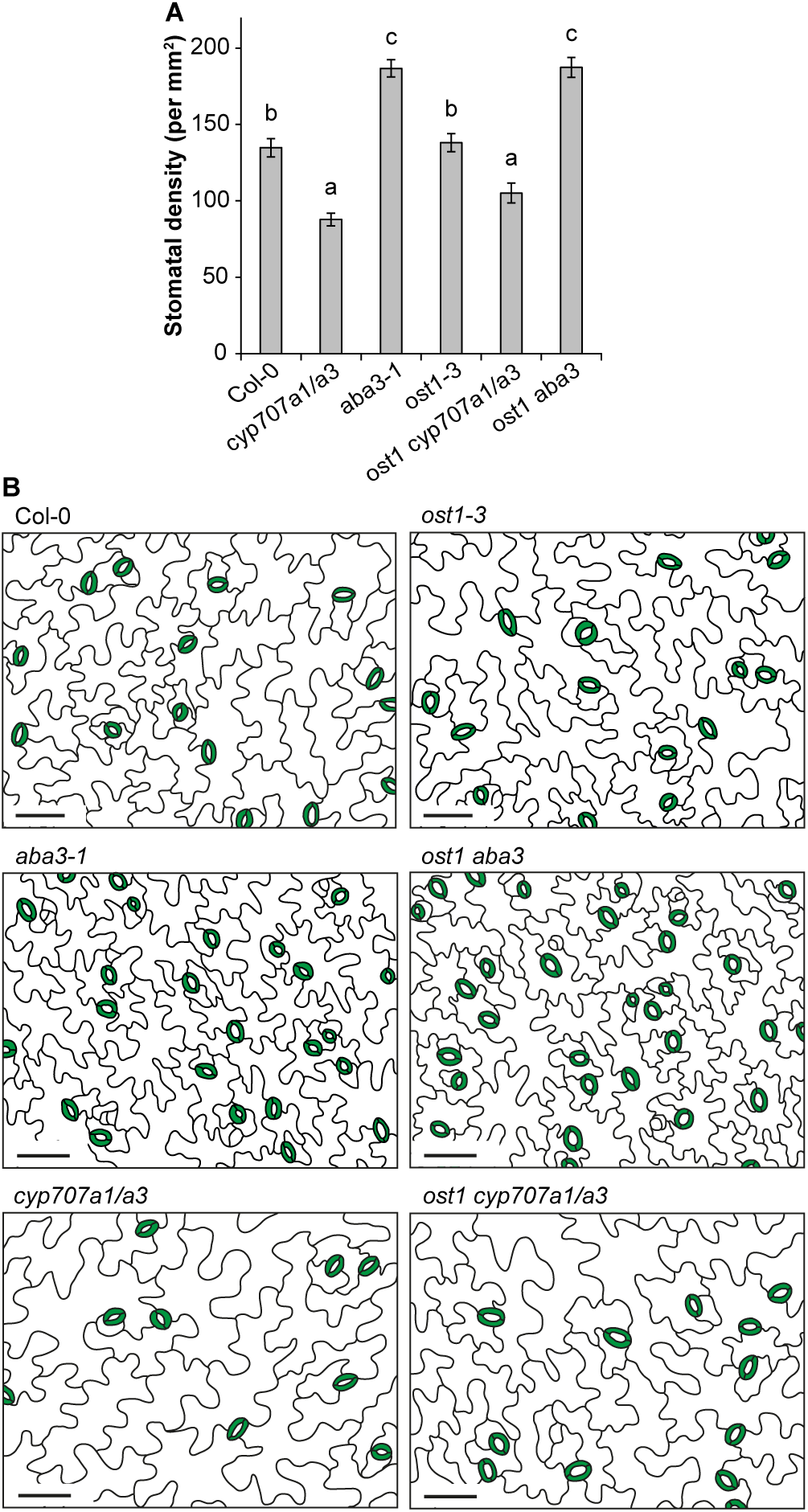
(A) Stomatal density of five-week-old plants. Letters denote statistically significant differences between lines (ANOVA with Tukey *post hoc* test, p < 0.05; n=24). (B) Tracing of epidermal impressions to illustrate the differences in stomatal densities between lines. The scale bar represents 50 μm.

To further characterize the role of ABA levels and OST1 in stomatal regulation, we tested the responses of single mutants, *ost1 aba3* and *ost1 cyp707a1/a3* to closure-inducing factors (Fig. 4 A-D). The closure induced by all stimuli was significantly impaired in *ost1-3*, whereas *aba3-1* plants showed wild-type-like closure or, in the case of reduced air humidity and ABA, even a hypersensitive response. In response to darkness, reduced air humidity and elevated CO_2_, the behavior of *ost1 aba3* and *ost1 cyp707a1/a3* mutants was not significantly different from *ost1-3* single mutant (Fig. 4 A-D, E-H). In response to ABA, *ost1 aba3* plants regained a small response that was larger compared to *ost1-3*, but reduced compared to wild-type (Fig. 4 D, H). Nevertheless, *ost1 aba3* and *ost1 cyp707a1/a3* were clearly impaired in rapid stomatal responses, supporting the critical role of OST1 in the regulation of stomatal aperture to sudden changes in the environment.

**Figure 4.**
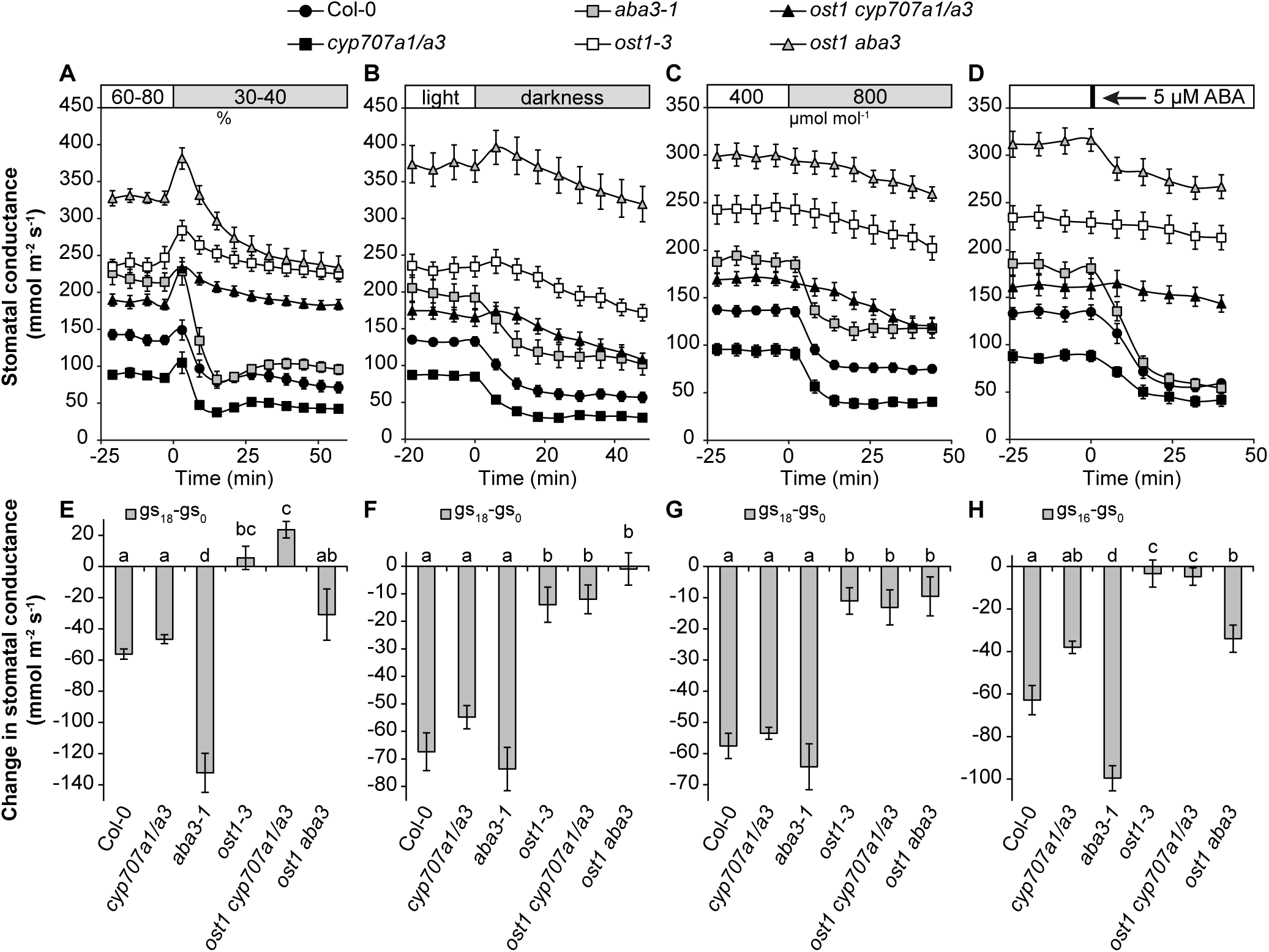
(A-D) Time courses of stomatal conductances in response to reduced air humidity (A), darkness (B), elevated CO_2_ (C) and ABA treatment (D). (E-H) Changes in stomatal conductance during the first 18 minutes (first 16 min in the ABA treatment). Letters denote statistically significant differences between lines (ANOVA with Tukey unequal N HSD *post hoc* test, p < 0.05; n=8-15).

## Discussion

Understanding stomatal function is critical for breeding plants with improved properties in water limiting conditions. As increased water loss from plants could be the result of either more open stomata or an increased number of stomata, the regulatory interplay between these traits is an important issue to be resolved. Changes in stomatal aperture width and density resulted in altered stomatal conductance, as suggested by our results (Fig. 2, Fig. 3 A). Through genetic manipulation of ABA levels, measurements of stomatal conductance, aperture, density and responses to various treatments, we propose that the overall water flux through stomata is the sum of two signaling pathways: an ABA-dependent pathway that is OST1-independent and regulates stomatal density; and an OST1-dependent pathway that regulates rapid changes of stomatal aperture. This conclusion is supported by our results which show significant differences in stomatal conductance due to differences in stomatal density and aperture between the single *ost1-3* mutant and its double and triple mutants where ABA levels are genetically reduced (*ost1 aba3*) and increased (*ost1 cyp707a1/a3*), respectively, while responses to various environmental stimuli are impaired in these mutants. It remains to be clarified whether other SnRKs besides OST1 (SnRK2.6) are involved in the ABA-dependent signaling involved in stomatal development. Alternatively, stomatal density could be determined by a SnRK-independent mechanism. However, the recent findings that a mutant lacking six ABA receptors and the PP2C mutants *abi1-1* and *abi2-1* also showed higher stomatal densities (Tanaka et al., 2013; Merilo et al., 2018) indicate that the canonical ABA signaling pathway starting with ABA receptors (Fig. 1) is involved in the regulation of stomatal development.

The change in stomatal conductance of mutants with altered concentrations of ABA appears to result from a change in stomatal density (Fig. 3A), aperture width or both, as in *aba3-1* (Fig. 2). The *aba3-1* mutant is relatively mildly impaired in ABA biosynthesis and still contains approximately 45% of wildtype ABA levels (Merilo et al., 2018). Mutants with more severely impaired ABA biosynthesis including *aba2-11* or *nced3 nced5* have considerably higher stomatal conductance than *aba3-1* (Merilo et al., 2018). Thus, the influence of ABA on stomatal aperture or density might become more prominent in plant lines where ABA concentrations are more severely reduced. By using stronger ABA-deficient lines including *aba2-11* or *nced3 nced5* (Merilo et al., 2018) or growing plants in water deficit conditions and measuring aperture and density may help to understand the balance between stomatal density and aperture in determining water flux through plants i.e. stomatal conductance. Stomatal density appeared to be more sensitive to reduced ABA levels than to increased levels as *cyp707a1/a3* showed only reduced density compared to wildtype (Fig. 2B). In order to breed crops for future climate, we need to understand the contribution of both stomatal density and stomatal aperture to plant water relations (see also Hughes et al., 2017). The latter is subjected to a rapid and up-to-date environmental control, whereas the former is fixed during plant development.

The triple mutant *snrk2.2 snrk2.3 snrk2.6* is completely impaired in ABA responses, including seed germination and gene expression (Fujii and Zhu, 2009; Umezawa et al., 2009). Thus, the OST1-independent mechanism regulating stomatal density could be genetically redundant among this group of SnRKs. Unfortunately the severe developmental defects of the *snrk* triple mutant (Fujii and Zhu, 2009) make it difficult to directly test this hypothesis. The *ost1* mutant was previously shown to completely lack stomatal responses to ABA (Mustilli et al., 2002), and applied at 5 µM, the *ost1* mutant is unresponsive to ABA (Fig. 4D). In an attempt to clarify if there is genetic redundancy among the SnRKs also in stomatal function, we treated *ost1-3* plants with very high 50 µM ABA, which induced a partial stomatal closure (Fig. S1). This supports earlier findings indicating that, besides OST1 there are other components, including SNRK2.2 and SNRK2.3 and other possible kinases, such as calcium dependent protein kinases (Brandt et al., 2015) or GHR1 (GUARD CELL HYDROGEN PEROXIDE_RESISTANT1) (Hua et al., 2012) acting in the ABA signaling pathway, that might contribute to ABA-induced stomatal closure. Other kinases may also explain the increased aperture width of *ost1 aba3* double mutant compared to single mutants. Several mitogen-activated protein kinases (MAPK) are involved in stomatal development (Wang et al., 2007) and ABA signaling (Jammes et al., 2009), indicating that MPKs might also contribute in the regulation of stomatal development and aperture in an ABA-dependent manner. Therefore, there are several candidate kinases in addition to OST1 in regulating stomatal responses to ABA.

Our results presented here show that it is possible to separate ABA signaling pathways that regulate stomatal aperture versus stomatal development. This information can be useful to breed separately for these traits to obtain plants suited for either rapidly changing environmental conditions or for conditions characterized by long-term drought.

## Acknowledgments

No conflict of interest declared.

This work was supported by the Estonian Ministry of Science and Education (IUT2-21 to H.K. and PUT-1133 to E.M.), the European Regional Development Fund (Center of Excellence in Molecular Cell Engineering CEMCE to H.K.), the Academy of Finland (grant number #307335, Center of Excellence in Molecular Biology of Primary Producers 2014-2019 to M.B.)

---

**Figure S1.**
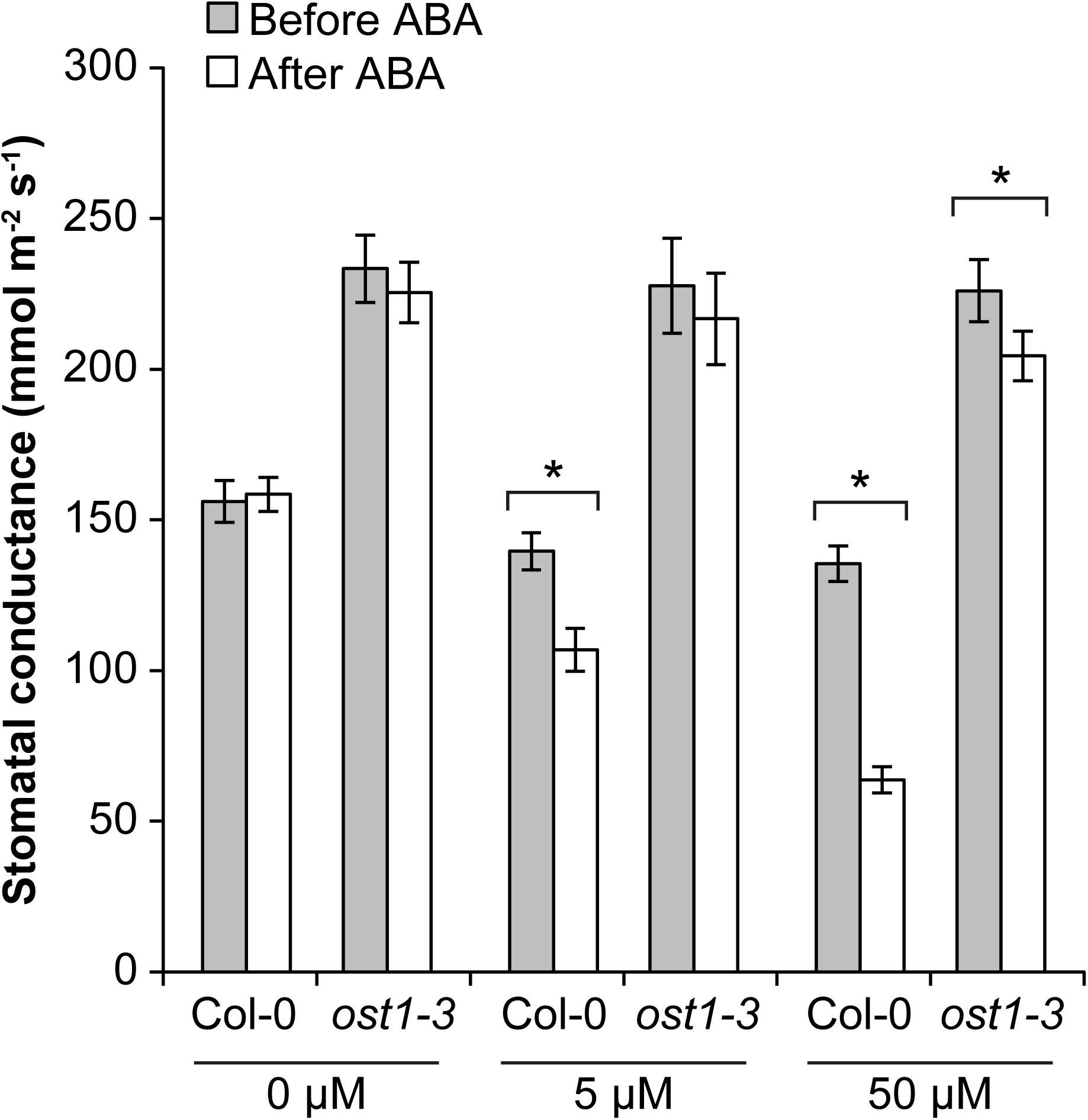
Stomatal response of Col-0 and *ost1-3* mutant to foliar ABA spraying (0 µM, 5 µM and 50 µM). Average (± SE, n=9) stomatal conductance before and 56 min after treatment with ABA. Statistically significant differences between post- and pretreatment stomatal conductance values are denoted by * (Repeated measures ANOVA with Tukey post hoc test, p<0.05).

